# Hiding in plain sight: rare *Bacillus cereus* group species are present among unassembled metagenomic samples in public databases

**DOI:** 10.64898/2026.10.02.756148

**Authors:** Vignesh Ramnath, Vishnu Raghuram, Laura M. Carroll

**Affiliations:** Department of Clinical Microbiology, Umeå University, Umeå, Sweden; Science for Life Laboratory (SciLifeLab), Umeå University, Umeå, Sweden; Laboratory for Molecular Infection Medicine Sweden (MIMS), Umeå University, Umeå, Sweden; Umeå Centre for Microbial Research (UCMR), Umeå University, Umeå, Sweden; Integrated Science Lab (IceLab), Umeå University, Umeå, Sweden

**Keywords:** Bacillus cereus group, *Bacillus manliponensis*, *Bacillus gaemokensis*, *Bacillus bingmayongensis*, *Bacillus clarus*, metagenomics

## Abstract

The *Bacillus cereus* group is a complex of closely related species, which vary in terms of their geographic spread, industrial utility, and pathogenic potential. In addition to encompassing well-known species represented by thousands of genomes (e.g., bioterrorism agent *B. anthracis*, foodborne pathogen *B. cereus*, biopesticide *B. thuringiensis*), the *B. cereus* group houses “rare” species discovered/first sequenced over a decade ago. Until recently, four rare species (*B. manliponensis*, *B. bingmayongensis, B. gaemokensis*, “*B. clarus*”) were each represented by a single unique genome; as such, virtually nothing is known about these species beyond their type strains. We hypothesized that additional representatives of rare *B. cereus* group species may already be present in public databases as yet-unassembled metagenomes. To this end, we used species-specific markers to identify 7 metagenomic samples in NCBI’s Sequence Read Archive, which harbored *B. manliponensis*, *B. bingmayongensis,* or *B. gaemokensis*. Metagenomic assembly and binning yielded an isolate-quality *B. manliponensis* metagenome-assembled genome (MAG) from avocado orchard soil in Israel, plus a partial *B. bingmayongensis* MAG from commercial spinach spiked with Shiga toxin-producing *Escherichia coli*. For rare species represented by >1 unique genome, we compared our novel MAGs to all available genomes and identified strain– and lineage-specific genomic elements (e.g., sphingomyelinase C and hemolysin BL homologs, variably present in *B. manliponensis* and *B. bingmayongensis*, respectively). Altogether, our study indicates that some rare *B. cereus* group species may be more common and geographically widespread than previously thought, and highlights the utility of culture-independent methods for recovering rare/under-isolated taxa.

**Impact Statement:** Individual *B. cereus* group strains vary in terms of their pathogenic potential and industrial utility. While some strains cause illness in humans and/or animals (e.g., anthrax, emetic intoxication, diarrheal toxicoinfection, non-gastrointestinal illnesses), other strains have important biotechnological applications (e.g., biocontrol, plant growth promotion, production of antimicrobials and other natural products). Because many virulence– and biotechnological-related phenotypes are strain-dependent (rather than species-dependent), phenotypes associated with an individual strain (e.g., a species type strain) cannot be extrapolated to an entire *B. cereus* group species. Within this context, rare *B. cereus* group species represent a major gap in our understanding of *B. cereus* group evolution, ecology, and pathogenicity, as until 2021, each rare species was represented solely by its type strain genome. Here, we aggregate all publicly available genomes/MAGs of rare *B. cereus* group species and supplement them with their first novel representatives since 2024. Our efforts bring the total number of unique *B. manliponensis* and *B. bingmayongensis* genomes/MAGs to >2, allowing us to conduct the first-ever phylogeographic survey and strain-level assessment of functional potential for each species. Overall, our study represents a critical first step toward understanding rare *B. cereus* group species beyond their type strains.

**Data Summary:** Metagenome-assembled genomes (MAGs) generated in this study are available in BTyperDB (https://btyperdb.btyper.app/) and NCBI, under NCBI BioProject accession PRJNA1528975 (for the novel *B. manliponensis* MAG: NCBI BioSample accession SAMN63123299, BTyperDB ID BTDB_2026-0001324.1; for the novel *B. bingmayongensis* MAG: NCBI BioSample accession SAMN63123300, BTyperDB ID BTDB_2026-0001323.1). All BTyperDB genomes used for *B. cereus* group pan-genome construction are available via Zenodo (https://doi.org/10.5281/zenodo.22960684). NCBI accessions for all publicly available genomes used in this study are available in the Supplementary Material (available via Zenodo: https://doi.org/10.5281/zenodo.21939590). All code for running software is described in the article, with additional figure construction code available via GitHub (https://github.com/lmc297/weird_bacillus). Interactive phylogenies for *B. manliponensis* and *B. bingmayongensis* are available via BTracker (https://nextstrain.org/community/vigzy77/BTracker/Bacillus-cereus-group/BTyper3-sp-manliponensis(Core-genome-ML) and https://nextstrain.org/community/vigzy77/BTracker/Bacillus-cereus-group/BTyper3-sp-bingmayongensis(Core-genome-ML), respectively).

## INTRODUCTION

The *Bacillus cereus* group (also known as *B. cereus sensu lato* or *B. cereus s.l.*) is a complex of closely related, Gram-positive, spore-forming species, which, collectively, are widespread in nature and occupy a diverse range of habitats (soil, in particular) [1–3]. While there is currently no universally accepted taxonomy for the *B. cereus* group [3], the species complex consists of 32 validly published species and six effective species according to the International Code of Nomenclature of Prokaryotes (ICNP) as of 28 June 2026 (referred to collectively hereafter as “published *B. cereus* group species”; see the **Supplementary Text** for methodological details regarding how this list was constructed) [4]: *B. albus [5]*, “*B. altaicus*” [6]*, B. anthracis [7]*, *B. arachidis [8], B. basilensis [9]*, *B. bingmayongensis* (proposed as a novel species in 2014, but recognized as validly published in 2026) [10, 11]*, B. bruguierae [12], B. cereus sensu stricto* (*s.s.*) [13], “*B. clarus*” [14]*, B. cytotoxicus [15]*, *B. dicomae [16]*, *B. fungorum [17]*, *B. gaemokensis* (proposed as a novel species in 2010, but recognized as validly published in 2021) [18], *B. hominis* (now considered a later heterotypic synonym of *B. mycoides*) *[19, 20]*, *B. luti [5]*, *B. manliponensis* (proposed as a novel species in 2011, but recognized as validly published in 2021) [21, 22], *B. mobilis [5]*, *B. multifaciens [23], B. mycoides [24]*, *B. nitratireducens [5]*, *B. pacificus [5]*, *B. paramobilis [19]*, *B. paramycoides [5]*, *B. paranthracis [5]*, “*B. pfraonensis*” [25], “*B. pretiosus*” [26]*, B. proteolyticus [5]*, *B. pseudomycoides [27]*, *B. rhizoplanae [28]*, *B. sanguinis [19]*, “*B. shihchuchen*” [29], “*B. silvicola*” [30]*, B. thuringiensis [31]*, *B. toyonensis* [32], *B. tropicus [5]*, *B. weihenstephanensis* (now considered a later heterotypic synonym of *B. mycoides*) *[33, 34]*, *B. wiedmannii [35]*, and *B. yunxiaonensis [12]*.

For each of the 38 published *B. cereus* group species, at least one representative (typically, the species type strain) has been characterized using whole-genome sequencing (WGS; per NCBI Datasets, https://www.ncbi.nlm.nih.gov/datasets/genome/, accessed 30 June 2026) [36]. However, beyond their type strain/species representative genomes, *B. cereus* group species vary wildly in terms of their representation in public genomic databases. Some species (e.g., bioterrorism agent *B. anthracis*, foodborne pathogen *B. cereus s.s.*, biopesticide *B. thuringiensis*) are relatively well-studied and well-sampled/sequenced (i.e., represented in public databases by hundreds-to-thousands of genomes each) [37, 38]. Comparatively, other species are represented by fewer than 5 genomes each, and sometimes, only by one unique genome from a single strain (i.e., the species type strain; as of 4 August 2026, “*B. clarus*”, *B. dicomae*, *B. gaemokensis, B. multifaciens*; **Supplementary Text**).

Predictably, most *B. cereus* group species represented by very few (<5 genomes) were discovered and published relatively recently (e.g., *B. bruguierae, B. multifaciens,* and *B. yunxiaonensis*, published as novel species in 2025; *B. dicomae* and *B. pfraonensis* in 2023; *B. sanguinis* in 2021; *B. fungorum* in 2020; **Supplementary Text**) [12, 16, 17, 19, 23, 25], indicating that continued sampling, isolation, and WGS efforts will likely uncover additional representatives of these species in the future. However, for several under-represented *B. cereus* group species, over a decade has passed since their discovery, with no-to-few novel genomes deposited in public databases, nor any novel isolates deposited in publicly accessible culture collections (referred to hereafter as “rare” *B. cereus* group species; **Figure 1**): (i) *B. manliponensis*, published in 2011 (via type strain BL4-6, isolated from oil-contaminated foreshore tidal flat sediment of the Yellow Sea, South Korea) [21]; (ii) *B. bingmayongensis*, published in 2014 (via type strain FJAT-13831, isolated in 2010 from a soil sample collected from the pit of Emperor Qin’s Terra-cotta Warriors in Shaanxi, China) [10]; (iii) *B. gaemokensis*, published in 2010 (via type strain BL3-6, isolated from tidal flat sediments of the Yellow Sea, South Korea) [18]; and (iv) “*B. clarus*”, first sequenced in 2014 and published as a novel species in 2020 (via type strain ATCC 21929, isolated from soil in Papua New Guinea). Notably, most rare *B. cereus* group species are relatively distantly related to all other species in the complex (for *B. manliponensis, B. gaemokensis,* and “*B. clarus*”, <88 average nucleotide identity [ANI] with all other species type strains; **Supplementary Text**) and represent unique lineages within the *B. cereus* group (**Figure 1A**, **Supplementary Figure S1**).

**Figure 1.**
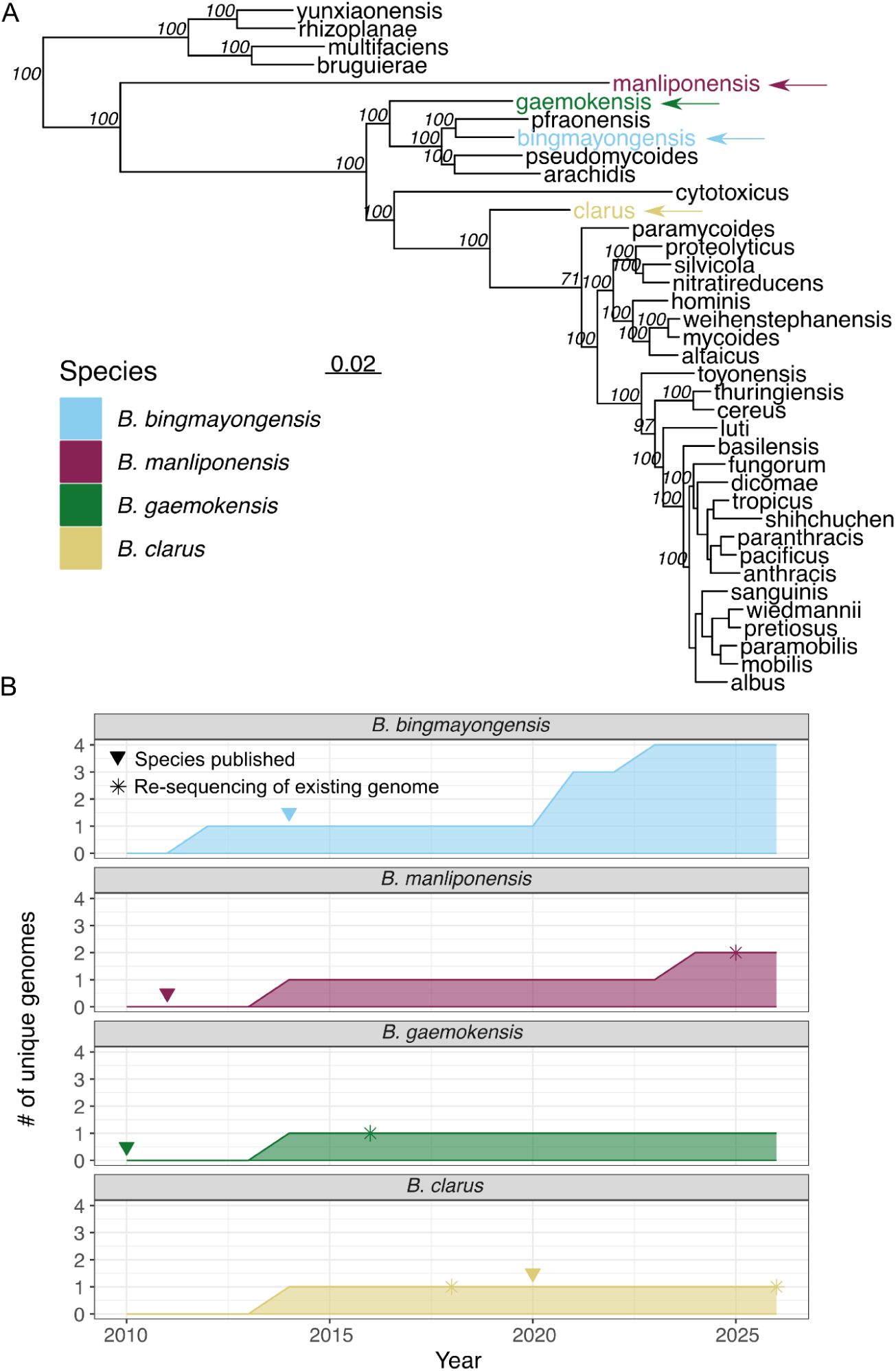
Despite being discovered and/or first sequenced over a decade ago, several *B. cereus* group species are represented in public databases by <5 (meta)genomes. (A) Rare species within the *B. cereus* group phylogeny. Rare species that were (i) discovered and/or first sequenced over a decade ago and (ii) are currently represented by <5 genomes/metagenome-assemble genomes (MAGs) in public databases are denoted by colored tip labels and arrows. The maximum likelihood phylogeny was constructed using the amino acid sequences of 1,270 single-copy orthogroups detected in all *B. cereus* group species type strains, plus an outgroup genome (i.e., *Ectobacillus panaciterrae* strain DSM 19096, NCBI Assembly accession GCF_000430785.1). Orthogroups were detected using OrthoFinder. The phylogeny is rooted at the outgroup (omitted for readability), with branch lengths reported in substitutions per site. Italicized branch labels denote ultrafast bootstrap support percentages (per IQ-TREE), with some labels omitted for readability (see **Supplementary Figure S1** for the unedited phylogeny). **(B)** Number of genomes/MAGs of unique rare *B. cereus* group species strains (Y-axis) available in public databases over time (in years; X-axis). Triangles denote the year each species was officially published, and asterisks denote the year a new genome/MAG of an existing strain was deposited in a public database (in all cases, this corresponded to re-sequencing of the species type strain).

Until 2021, the four rare *B. cereus* group species (i.e., *B. manliponensis*, *B. bingmayongensis*, *B. gaemokensis*, and “*B. clarus*”) were represented solely by their type strain genomes (**Figure 1B**). For *B. gaemokensis* and “*B. clarus*”, the type strain continues to be the sole species representative (**Figure 1B**). For *B. manliponensis* and *B. bingmayongensis*, genomes from additional strains have been published (*n* = 1 and 3 additional unique genome[s] since 2021, respectively; **Figure 1B**), but no evolutionary or comparative genomic analyses have been performed for either species. Consequently, virtually nothing is known about the rare *B. cereus* group species beyond their type strains. This knowledge gap is extremely problematic, as individual *B. cereus* group strains have long been known to vary in terms of their pathogenic potential and industrial utility/relevance [1]. Specifically, many virulence– and biotechnological-related phenotypes are strain-dependent rather than species-dependent (e.g., ability to cause anthrax/anthrax-like illness or foodborne emetic intoxication; cytotoxicity; ability to produce insecticidal crystal proteins) [1, 37, 39]. As such, phenotypes associated with an individual strain (e.g., a species type strain) cannot be extrapolated to an entire *B. cereus* group species. Taken together, rare *B. cereus* group species represent a major gap in our understanding of *B. cereus* group evolution, ecology, and pathogenicity, and thus hinder our ability to conduct accurate strain-level risk assessments.

Rare *B. cereus* group species have largely evaded global isolation and WGS efforts. However, in recent years, shotgun metagenomic sequencing has become cheaper, more accessible, and more globally widespread [40]. With the rapidly increasing amount of shotgun metagenomic sequencing data available in public databases, we hypothesized that global metagenomic sequencing efforts may have already captured rare *B. cereus* group species. To test this hypothesis, we leverage >6,000 *B. cereus* group genomes to identify “signatures” (i.e., genetic markers) of rare *B. cereus* group species. We then search NCBI’s Sequence Read Archive (SRA) for these signatures, with the goal of identifying previously unassembled metagenomes, which harbor rare *B. cereus* group species. Beyond providing a workflow for rapidly identifying rare species in unassembled metagenomes in SRA, our study represents a first attempt at understanding the evolution, ecology and functional potential of rare *B. cereus* group species.

## METHODS

### *B. cereus* group species type strain phylogeny construction

*B. cereus* group species type strain genomes (*n* = 38; **Supplementary Table S1**, **Supplementary Text**) and an outgroup genome (i.e., *Ectobacillus panaciterrae* strain DSM 19096, NCBI Assembly accession GCF_000430785.1) [41] were supplied as input to Prokka v1.14.6 [42] (parameters: ‘––kingdom Bacteria ––cpus 4 ––addgenes ––centre ––compliant’). The resulting protein FASTA files (extension ‘.faà) were supplied as input to OrthoFinder v3.1.5 [43–46], which was used to construct an alignment of single-copy orthogroups detected in all 39 genomes using the multiple sequence alignment method for gene tree inference, MAFFT as the aligner, RAxML-NG for tree construction, and 28 threads (parameters: ‘-t 28 –a 28 –M msa –A mafft –T raxml-ng’) [47, 48]. The resulting alignment (filename ‘SpeciesTreeAlignment.fà) was supplied as input to IQ-TREE v2.2.5 [49] and used to construct a maximum likelihood phylogeny using the optimal amino acid (AA) substitution model selected using ModelFinder (i.e., the ‘Q.plant+F+I+R8’ model) [50–54], one thousand replicates of the ultrafast bootstrap approximation [55], and 28 threads (parameters: ‘-m MFP –B 1000 –nt 28’). The resulting phylogeny was rooted at the outgroup (using the ‘root’ function from the ape v5.8-1 R package) [56] and annotated in R v4.6.1 [57] using the ggtree v4.2.0 package [58] (**Figure 1A, Supplementary Figure S1**).

### Identification of species-specific markers in publicly available metagenomic samples

All 6,013 *B. cereus* group genomes available in BTyperDB (accessed August 2022) were downloaded and annotated using bakta v1.8.2 (default parameters; database version: 10.5281/zenodo.7669534) [59]. GFF files produced by bakta were then supplied as input to Panaroo v1.5.1 for pan-genome analysis (parameters: ‘––clean-mode strict ––remove-invalid-genes –f 0.5 –a pan ––core_threshold 0.95 ––refind_prop_match 0.8 ––refind-mode strict’) [60]. The resulting Panaroo gene presence/absence matrix was then used to identify orthologous genes that were exclusive to each of *B. manliponensis* (NCBI Assembly accession GCA_000712595.1), *B. bingmayongensis* (NCBI Assembly accessions GCA_000299035.1 and GCA_019753385.1), *B. gaemokensis* (NCBI Assembly accession GCA_000712615.1), and “*B. clarus*” (NCBI Assembly accession GCA_000746925.1). This approach resulted in a total of 245 orthologous genes specific to *B. manliponensis*, 195 to *B. bingmayongensis*, 169 to *B. gaemokensis*, and 225 to “*B. clarus*” (referred to hereafter as “species-specific genes”; **Supplementary Datasets S1-S4**).

The nucleotide (DNA) sequence of each resulting species-specific gene (from Panaroo’s ‘pan_genome_reference.fà file) was supplied as a query (in FASTA format) to Pebblescout (https://pebblescout.ncbi.nlm.nih.gov, accessed 3 July 2026) [61] via the API (Application Programming Interface); Pebblescout was used to search both metagenomic databases, with “report type” set to “2” (i.e., to produce a summary report; remaining parameters were set to their defaults): (i) database = ‘metà, “Metagenomic, Volume 1”, i.e., all public metagenomic and metatranscriptomic runs in SRA before the end of 2021; (ii) database = ‘meta_vol2’, “Metagenomic, Volume 2”, i.e., all public metagenomic and metatranscriptomic runs in SRA before the end of 2023 except those present in “Metagenomic, Volume 1”. Each search (one for each species-specific gene/database combination) resulted in a list of samples (i.e., SRA Runs) in which a species-specific gene was detected (**Supplementary Tables S2-S5**).

In addition to species-specific genes, we used Pebblescout to search for species-specific alleles of *panC* (pantoate-β-alanine ligase), a *B. cereus* group core gene that is popular for assignment of phylogenetic groups (roughly equivalent to species) [38, 62, 63]. Briefly, *panC* alleles of *B. manliponensis, B. bingmayongensis*, *B. gaemokensis,* and “*B. clarus*” (identified in a previous study) [38] were acquired from BTyper3 v3.4.0 (https://github.com/lmc297/BTyper3/blob/master/btyper3/seq_panC_db/panC.fna). The nucleotide sequence of each *panC* allele was then searched against NCBI’s ‘metà and ‘meta_vol2’ databases using Pebblescout as described above. Each search (one for each *panC* allele/database combination) resulted in a list of samples (i.e., SRA Runs) in which a rare *B. cereus* group species *panC* allele was detected (**Supplementary Tables S6-S8**).

### Acquisition of metagenomic samples putatively harboring rare *B. cereus* group species

Using Pebblescout results for species-specific (i) genes and (ii) *panC* alleles (see section “Identification of species-specific markers in publicly available metagenomic samples” above), metagenomic samples meeting the following conditions were downloaded by supplying the corresponding SRA Run accession number to the ‘prefetch’ command in NCBI’s SRA Toolkit v3.0.8 (https://github.com/ncbi/sra-tools): (i) a rare *B. cereus* group species *panC* allele was detected in the sample with ≥80% coverage (per Pebblescout), and/or (ii) ≥15% of the species-specific genes were detected in the sample with ≥80% coverage (per Pebblescout; **Supplementary Tables S2-S8**). Each resulting SRA archive was supplied as input to the ‘fastq-dump’ command from SRA Toolkit (parameters: ‘––split-files ––gzip’), resulting in gzipped FASTQ file(s) with raw sequencing reads for each sample. One sample (SRA Run accession ERR1992808) was reportedly sequenced using Illumina paired-end reads, but produced only a single gzipped FASTQ file, and was treated as a single-end sample as a result. Two reportedly paired-end samples (SRA Run accessions SRR7186533 and SRR7186537) were split into forward– and reverse-read files by ‘fastq-dump’, but when undergoing taxonomic profiling, read pairs could not be discerned (see section “Taxonomic profiling” below). As such, these two samples were treated as single-end samples for taxonomic profiling, and as both paired-end and single-end samples for MAG construction (i.e., both samples underwent MAG construction twice, first as paired-end samples and second as single-end samples; see section “Construction and quality control of metagenome-assembled genomes” below).

### Taxonomic profiling

To assess the relative abundance of species within each metagenomic sample (see section “Acquisition of metagenomic samples putatively harboring rare *B. cereus* group species” above), taxonomic profiling was performed. Briefly, FastQC v0.12.1 (https://www.bioinformatics.babraham.ac.uk/projects/fastqc/) and MultiQC v1.16 [64] were used to assess the quality of all raw sequencing reads (default settings). Read trimming/filtering was then performed via fastp v0.23.4 [65], using the following settings (remaining settings were set to their defaults): ‘––length_required 36’ (all samples), ‘––detect_adapter_for_pè (paired-end samples only). FastQC and MultiQC were then used as described above to assess the quality of the resulting trimmed paired-end reads. Trimmed paired-end reads were then supplied as input to the mOTUs profiler v4.0.4 [66] and used to obtain the relative abundance of mOTUs (marker gene-based, species-level operational taxonomic units) in each metagenomic sample (using default settings and the mOTUs marker gene database, downloaded via the ‘motus downloadMGDB’ command, accessed 15 July 2026; **Supplementary Tables S9-S15**). Results were plotted in R using ggplot2 v4.0.3 [67].

### Construction and quality control of metagenome-assembled genomes

For each metagenomic sample (see section “Acquisition of metagenomic samples putatively harboring rare *B. cereus* group species” above), metagenome-assembled genomes (MAGs) were constructed using the nf-core/mag pipeline v5.4.2-g5dabb01 [68] and Nextflow v25.10.0 [69]. Briefly, for each sample, a comma-separated samplesheet was created using sample metadata from SRA (e.g., sequencing platform). Each samplesheet was then supplied as input to the ‘nextflow run nf-core/mag’ command (via the ‘––input’ parameter), with the following parameters (remaining parameters were set to their defaults): ‘––skip_prokkà (to skip MAG annotation with Prokka); ‘––skip_gtdbtk’ (to skip taxonomic assignment of MAGs using GTDB-Tk). For the three single-end samples (ERR1992808, SRR7186533, SRR7186537), the ‘––single_end’ parameter was additionally included. For four samples (NCBI SRA Run accessions SRR7186533, SRR7186537, ERR1992808, and ERR2185279), ALE [70] could not be executed successfully due to memory constraints; as such, for these samples, the ‘––skip_alè parameter was additionally included. Similarly, COMEBin v1.0.4 [71] could not be executed successfully for two samples due to memory/time constraints (NCBI SRA Run accessions ERR1992808 and ERR2185279), and the ‘––skip_comebin’ option was additionally included.

nf-core/mag was then used to generate MAGs for each sample, testing a range of assembly and binning methods in the process. Major steps in the overall workflow were as follows (all steps used nf-core/mag default settings): (i) assessment of raw read quality via FastQC v0.12.1 (https://www.bioinformatics.babraham.ac.uk/projects/fastqc/); (ii) read trimming with fastp v1.0.1 [65]; (iii) PhiX read removal with Bowtie2 v2.4.2 [72]; (iv) assessment of trimmed read quality via FastQC; (v) *de novo* assembly with both MEGAHIT v1.2.9 [73] and metaSPAdes v4.1.0 [74]; (vi) assessment of *de novo* assembly quality with MetaQUAST v5.0.2 [75] (all samples) and ALE v20180904 [70] (all but the samples specified above); (vii) gene identification with Prodigal v2.6.3 [76]; (viii) alignment of reads to metagenomic contigs with Bowtie2; (ix) binning of metagenomic contigs into MAGs with COMEBin v1.0.4 (all but the samples listed above) [71], CONCOCT [77], MaxBin2 v2.2.7 [78], MetaBAT2 v2.17 (v2.15 for the nf-core/mag “METABAT2_JGISUMMARIZEBAMCONTIGDEPTHS_SHORTREAD” process) [79], MetaBinner v1.4.4-0 [80], and SemiBin2 v2.2.0 [81]; (x) evaluation of MAG (i.e., bin) quality with MetaQUAST and BUSCO v6.0.0 [82].

### Identification of rare *B. cereus* group species MAGs

To identify putative *B. cereus* group MAGs, the ‘skani dist’ command from skani v0.1.4 [83] was used to calculate average nucleotide identity (ANI) values between MAGs (reference genome; see section “Construction and quality control of metagenome-assembled genomes” above) and the type strain genome of the corresponding rare *B. cereus* group species (query; i.e., NCBI Assembly accession GCF_000712595.1 for *B. manliponensis*, GCF_000299035.1 for *B. bingmayongensis*, and GCF_001590835.1 for *B. gaemokensis*); for moderately and highly fragmented MAGs (i.e., MetaQUAST N50 <10 Kb and <3 Kb, respectively), the ‘––medium’ and ‘––slow’ parameters were additionally included, respectively (as suggested by skani). For metagenomic samples that putatively contained more than one rare *B. cereus* group species (i.e., per the mOTUs profiler; see section “Taxonomic profiling” above), skani calculations were repeated using each rare species query genome. MAGs that produced a valid ANI value (regardless of the value) were treated as putative *B. cereus* group MAGs and underwent additional quality control via CheckM v1.2.2 [84] (’checkm lineage_wf’ with default parameters except ‘-t 28 –f checkm_out.txt ––tab_tablè) and taxonomic classification using GTDB-Tk v2.6.1 (’gtdbtk classify_wf’ with default settings, 28 CPUs, and GTDB release r226) [85] and BTyper3 v3.4.0 (default settings) [38]. MAGs that belonged to a rare *B. cereus* group species using GTDB-Tk, BTyper3’s ANI-based genomospecies assignment method, and/or BTyper3’s *panC* Group assignment method were considered to be rare species MAGs and were used in subsequent steps (**Supplementary Tables S16-S18)**.

### Comparison of novel MAGs to publicly available genomes/MAGs

For each rare *B. cereus* group species represented by >1 unique genome (i.e., *B. manliponensis*, *B. bingmayongensis*), the highest-quality novel MAG(s) (i.e., MAGs generated in this study; see “Results” below) were compared to all publicly available genomes and MAGs of the respective species (all URLs accessed 12 July 2026 for *B. manliponensis* and 19 July 2026 for *B. bingmayongensis*). Publicly available genomes/MAGs were identified by first querying each species name in the Genome database of NCBI Datasets (https://www.ncbi.nlm.nih.gov/datasets/genome/) [36, 86]; all GenBank genomes that resulted from the search were downloaded in FASTA format. Second, to ensure that rare species genomes classified in NCBI under alternative taxonomic IDs were not excluded, each species name was queried in GTDB (https://gtdb.ecogenomic.org/) [87], and any additional genomes classified as the respective rare species via GTDB were downloaded from NCBI. Finally, to ensure that previously assembled MAGs not included in NCBI were not excluded, each species name was queried in the mOTUs (https://motus-db.org/home) [88] and SPIRE (https://spire.embl.de/) [89] databases.

Altogether, this search resulted in 3 publicly available genomes for *B. manliponensis* (all from searching NCBI) and 4 publicly available genomes for *B. bingmayongensis* (3 from searching NCBI and 1 additional genome from searching GTDB; **Supplementary Tables S19-S20**). For *B. gaemokensis* and “*B. clarus*”, this search resulted in one unique genome each (i.e., derived from the species type strain; **Figure 1B**); this, combined with the fact that our MAG construction efforts did not yield any novel *B. gaemokensis* or “*B. clarus*” MAGs, meant that we were unable to query *B. gaemokensis* and “*B. clarus*” in subsequent steps.

For each rare *B. cereus* group species represented by >1 unique genome (i.e., *B. manliponensis*, *B. bingmayongensis*), all genomes/MAGs underwent the following analyses: (i) quality control using CheckM (see section “Identification of rare *B. cereus* group species MAGs” above) and QUAST (v5.2.0 with ‘––min-contig 1’) [90]; (ii) taxonomic confirmation using GTDB-Tk and BTyper3 (see section “Identification of rare *B. cereus* group species MAGs” above); (iii) annotation using Prokka v1.14.6 (parameters: ‘––kingdom Bacteria ––cpus 4 ––addgenes ––centre ––compliant’; **Supplementary Tables S19-S20**) [42]. GFF files produced by Prokka were supplied as input to Panaroo v1.3.4 [60] for pan-genome analysis (parameters: ‘––clean-mode strict –a core ––aligner mafft ––core_threshold 0.95 –t 28 ––remove-invalid-genes –f 0.5’; **Supplementary Tables S21-S22**) [47]. Orthologous genes identified by Panaroo, which were exclusively present in or absent from the novel MAGs generated here were annotated using eggNOG-mapper v2.1.13 [91] (accessed via the Galaxy Europe server, https://usegalaxy.eu/login/start, and executed with the following parameters, with the remaining set to their defaults: eggNOG database v5.0.2, ‘Set of GO terms that should be used for annotation’ set to ‘All’, ‘Type of sequences’ set to ‘CDS’; **Supplementary Tables S23-S26**) [92–94]. Exclusively present genes were additionally supplied to NCBI’s BLAST webserver (https://blast.ncbi.nlm.nih.gov/Blast.cgi, accessed 21 July 2026 for *B. manliponensis* and 22 July 2026 for *B. bingmayongensis*) for annotation with BLASTX (default parameters, plus genetic code set to ‘Bacteria and Archaea (11)’ and database set to ‘Non-redundant protein sequences (nr)’; **Supplementary Datasets S5-S6**) [95]. The ‘vegdist’ function from the vegan v2.7-5 R package [96] was additionally used to calculate Jaccard distances between genomes/MAGs within each species, using the Panaroo gene presence/absence matrix (’gene_presence_absence.Rtab’) as input and ‘method = “jaccard“, binary = T’; results were plotted in R using the ‘pheatmap’ function from pheatmap v1.0.13 [97].

The filtered core gene alignment produced by Panaroo (’core_gene_alignment_filtered.aln’) was additionally supplied as input to IQ-TREE and used to construct an unrooted maximum-likelihood phylogeny as described above (see section “*B. cereus* group species type strain phylogeny construction”), with the “GTR+F+G4” and “GTR+F+I” nucleotide substitution models selected by ModelFinder as the best-fit models for *B. manliponensis* and *B. bingmayongensis*, respectively (**Supplementary Datasets S7-S8**) [98, 99]. To construct a rooted maximum-likelihood phylogeny for each species, Panaroo was run a second time as described above, but with the addition of an outgroup genome (for both *B. manliponensis* and *B. bingmayongensis*, *B. pseudomycoides* str. DSM 12442, NCBI RefSeq Assembly accession GCF_000161455.1; GFF file constructed using Prokka as described above). The resulting filtered core gene alignment was supplied to IQ-TREE and used to construct a maximum likelihood phylogeny as described above, with the “GTR+F+I” nucleotide substitution model selected as the best-fit model for *B. manliponensis*, and the “GTR+F+I+G4” model for *B. bingmayongensis* (**Supplementary Datasets S9-S10**). Phylogenies were plotted alongside metadata in R using the ggtree and ape packages.

Pairwise ANI values were calculated between all genomes/MAGs within each rare species using skani as described above (see section “Identification of rare *B. cereus* group species MAGs”; **Supplementary Tables S27-S28**). For each novel MAG sharing a high degree of genomic similarity with publicly available genomes/MAGs (≥99.0 ANI), Snippy v4.6.0 (https://github.com/tseemann/snippy; default settings) [100–106] was used to identify core SNPs in the novel MAG, using the GenBank file of the publicly available genome/MAG as a reference (downloaded from NCBI’s FTP site; **Supplementary Table S29**). Heatmaps of ANI dissimilarities (100-ANI) were constructed in R using the pheatmap package.

### *B. cereus* group virulence factor detection

Many important *B. cereus* group virulence factors (e.g., anthrax toxin and capsule genes; the cereulide synthetase biosynthetic gene cluster) are harbored on plasmids [1]. Therefore, in addition to detecting virulence factors in all rare species genomes/MAGs at the amino acid (AA) level (using BTyper3; see section “Comparison of novel MAGs to publicly available genomes/MAGs” above), we used BTyper3’s implementation of translated nucleotide BLAST (tblastn) [107] to detect virulence factors in all assembled contigs produced for each metagenomic sample at the AA level (default settings; i.e., to detect virulence factors harbored on plasmids, which did not get binned with rare species MAGs). Some *B. cereus* group virulence factors can be detected with high coverage but relatively low AA identity (<70%; e.g., non-hemolytic toxin Nhe) [38, 108]. Thus, to avoid using hard AA identity and coverage cutoffs, all raw tblastn results were manually inspected to assess virulence factor presence/absence (**Supplementary Tables S30-S45**).

### Antimicrobial resistance determinant detection

All rare *B. cereus* group genomes/MAGs underwent antimicrobial resistance (AMR) determinant detection using AMRFinderPlus v3.12.8 [109]. Briefly, contig FASTA (’.fnà), protein FASTA (’.faà), and GFF (’.gff’) files produced by Prokka (see section “Comparison of novel MAGs to publicly available genomes/MAGs” above) were supplied as input to the ‘-n’, ‘-p’, and ‘-g’ options of ‘amrfinder’, respectively; ‘amrfinder’ was executed using the following parameters (remaining were left to their defaults) ‘––annotation_format prokka ––plus’, database v2024-01-31.1 (**Supplementary Tables S46-47**).

### Identification of mobile genetic elements

Detection of mobile genetic elements (MGEs) within each rare *B. cereus* group genome/MAG (see section “Comparison of novel MAGs to publicly available genomes/MAGs” above) was performed using the ‘genomad end-to-end’ command from geNomad 1.12.0 [110] with geNomad database v1.9 and the following parameters (remaining parameters were set to their defaults): ‘––conservative ––cleanup ––enable-score-calibration ––lenient-taxonomy ––full-ictv-lineagè (**Supplementary Tables S48-S51**). To compare rare *B. cereus* group viral sequences to previously observed prophage, viral sequences identified by geNomad were supplied as input to Prokka (parameters: ‘––kingdom Viruses ––addgenes ––centre ––compliant’), and the resulting GenBank files were concatenated alongside all prophage GenBank files from Prophage-DB (processed as described previously) [111, 112]. IGUA v0.2.1 [112] was used to cluster all viral sequences into gene cluster families (GCFs) based on their protein composition (’-j 72’ and default settings; **Supplementary Tables S52-S53**). Viral sequences within select GCFs were compared and visualized using clinker v0.0.31 (parameters: ‘-j 1 –dl “,” –dc 4’) [113].

One unique viral sequence with a direct terminal repeat (DTR) topology (discussed in detail below in the “Results” section) was further investigated by submitting the amino acid (AA) sequence of each protein within the predicted virus to the NCBI BLAST webserver (https://blast.ncbi.nlm.nih.gov/), using the following command from Biopython v1.86 [114]: ‘NCBIWWW.qblast’(“blastp”, “nr”, record.format(“fasta”), format_type=“Text”)’.

### Biosynthetic gene cluster detection and characterization

Biosynthetic gene clusters (BGCs) were detected in each rare *B. cereus* group genome/MAG (see section “Comparison of novel MAGs to publicly available genomes/MAGs” above) using (i) GECCO v0.9.8 (parameters for ‘gecco run’: ‘-j 1 ––mask –m 0.3’) [115] and (ii) antiSMASH v8.0.4 (parameters: ‘––taxon bacteria –c 4 ––cc-mibig ––cb-knownclusters ––cb-general ––tigrfam ––pfam2go ––genefinding-tool prodigal-m’) [116]. The resulting BGCs (in GenBank format) were concatenated alongside previously characterized BGCs from the MIBiG v4.0 database [117], clustered into GCFs (gene cluster families) using IGUA (’-j 8’ and default settings; **Supplementary Tables S54-S55**), and compared/visualized using clinker (parameters: ‘-j 1 –dl “,” –dc 4’).

## RESULTS

### Genomic markers of rare *B. cereus* group species are present in publicly available, unassembled metagenomic samples

Using data from our large-scale genomic studies of the *B. cereus* group [38, 59], we identified species-specific markers, which were potentially capable of distinguishing rare *B. cereus* group species from other members of the species complex, including: (i) 834 total species-specific genes, which were uniquely found in rare *B. cereus* group species (i.e., 245 genes specific to *B. manliponensis*, 195 to *B. bingmayongensis*, 169 to *B. gaemokensis*, and 225 to “*B. clarus*”); and (ii) alleles of the *B. cereus* group core gene *panC*, which were unique to each rare species (see “Methods” above for details). We then used Pebblescout to search all metagenomic samples in NCBI’s SRA database for each species-specific marker, with the goal of identifying yet-unassembled metagenomic samples, which putatively harbored rare *B. cereus* group species (**Figure 2A**).

**Figure 2.**
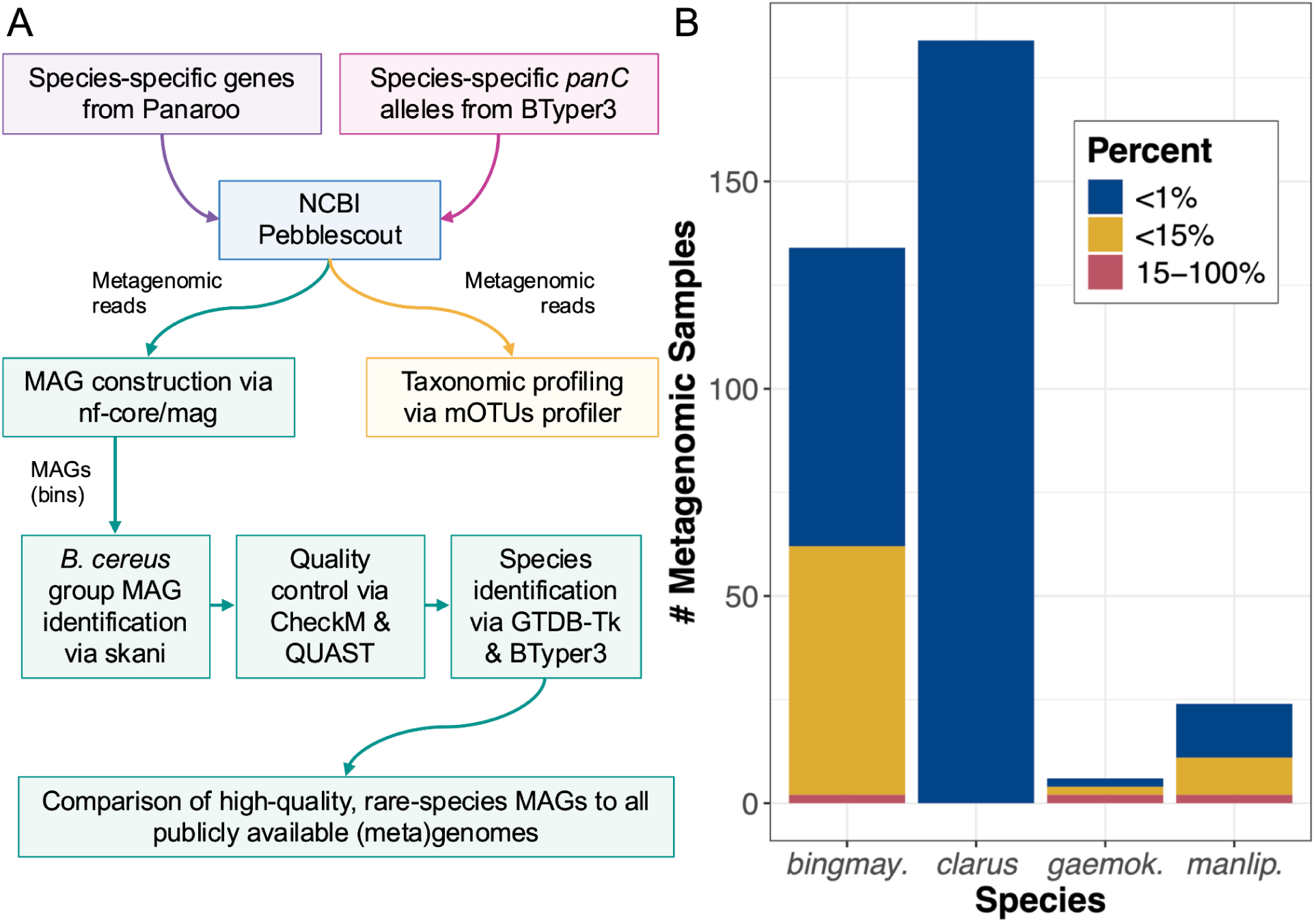
Markers of rare *B. cereus* group species are present in hundreds of publicly available, yet-unassembled metagenomes. **(A)** Bioinformatic workflow used to identify/characterize publicly available metagenomes harboring rare *B. cereus* group species. MAG, metagenome-assembled genome. **(B)** Number (#) of metagenomic samples (i.e., Sequence Read Archive [SRA] Runs; Y-axis) with ≥1 rare *B. cereus* group species-specific gene (detected with ≥80% coverage via Pebblescount; X-axis). Bars are colored by the number of samples harboring a given percentage of species-specific genes (out of 245 species-specific genes for *B. manliponensis*, 195 for *B. bingmayongensis*, 169 for *B. gaemokensis*, and 225 for “*B. clarus*”).

Using a cutoff of ≥80% coverage (per Pebblescout), our search identified 331 metagenomic samples in SRA (i.e., SRA Runs), which harbored at least one rare *B. cereus* group species-specific gene (**Figure 2B**). *B. gaemokensis*-specific genes were detected in the fewest number of samples (*n* = 6); however, two of these samples (33.3%) possessed a relatively large number of *B. gaemokensis*-specific genes (35/169 *B. gaemokensis*-specific genes, 20.7%; **Figure 2B, Supplementary Table S4**). While “*B. clarus*”-specific genes were detected in a relatively large number of samples (*n* = 184, the most of any rare *B. cereus* group species), only 1-2 “*B. clarus*”-specific genes were detected per sample (out of 225 “*B. clarus*”-specific genes, 0.44-0.89%; **Figure 2B**), suggesting that these may be spurious hits.

Compared to species-specific genes, the presence of species-specific *panC* alleles appeared to be a much more conservative metric of rare *B. cereus* group species presence, as only five metagenomic samples harbored rare *B. cereus* group species *panC* alleles (*n* = 2, 1, and 2 samples harboring *B. manliponensis, B. bingmayongensis,* and *B. gaemokensis panC* alleles, respectively, with ≥80% coverage per Pebblescout; **Supplementary Tables S6-S8**). Notably, the “*B. clarus*” *panC* allele could not be detected in any metagenomic samples from SRA with ≥80% coverage. The absence of the “*B. clarus*” *panC* allele, combined with the low prevalence of samples harboring multiple “*B. clarus*”-specific genes, provided further evidence that a subset of “*B. clarus*”-specific genes could produce spurious hits (**Supplementary Table S5**).

Taking into account both species-specific genes and *panC* alleles, we prioritized metagenomic samples that met the following conditions: (i) a rare *B. cereus* group species *panC* allele was detected in the sample with ≥80% coverage (per Pebblescout; *n* = 5 samples), and/or (ii) ≥15% of the species-specific genes were detected in the sample with ≥80% coverage (per Pebblescout, *n* = 6 samples; see “Methods” section above). Based on these criteria, 7 metagenomic samples were selected to undergo taxonomic profiling and MAG construction (*n* = 2, 3, and 2 samples putatively harboring *B. manliponensis*, *B. bingmayongensis*, and *B. gaemokensis*, respectively). We discuss our taxonomic profiling and MAG construction efforts for each detected rare *B. cereus* group species below, and for species with >1 unique genome/MAG available (i.e., *B. manliponensis, B. bingmayongensis*), we perform the first in-depth, species-wide comparative genomic analysis and assessment of functional potential.

### *B. manliponensis* is present in freshwater-irrigated soil from an Israeli avocado orchard

Complete/nearly complete *B. manliponensis panC* alleles were detected in two metagenomic samples from SRA (Pebblescout coverage 100.0% and 92.1% for NCBI SRA Run accessions SRR12632054 and SRR12632052, respectively; **Supplementary Table S6**). Both samples were biological replicates, derived from freshwater-irrigated soil from the same avocado orchard (“orchard 2”) in Israel, collected on the same date (15 September 2019) [118]. In addition to harboring (near-)complete *B. manliponensis panC* alleles, both samples possessed nearly all *B. manliponensis*-specific genes (216/245 genes, 88.2% for SRR12632054, and 197/245 genes, 80.4% for SRR12632052; **Figure 2B**, **Supplementary Table S2**). Using the mOTUs profiler, *B. manliponensis* was detected in both soil samples with a relative abundance of <1% (**Figure 3**, **Supplementary Figure S2**, **Supplementary Tables S9-S10**). Together, these results indicated that it may be possible to obtain a high-quality *B. manliponensis* MAG from at least one of these samples. To this end, we constructed MAGs for each soil sample, testing all combinations of two *de novo* assembly methods (MEGAHIT and metaSPAdes) and six binning methods (COMEBin, CONCOCT, MaxBin2, MetaBAT2, MetaBinner, and SemiBin2).

**Figure 3.**
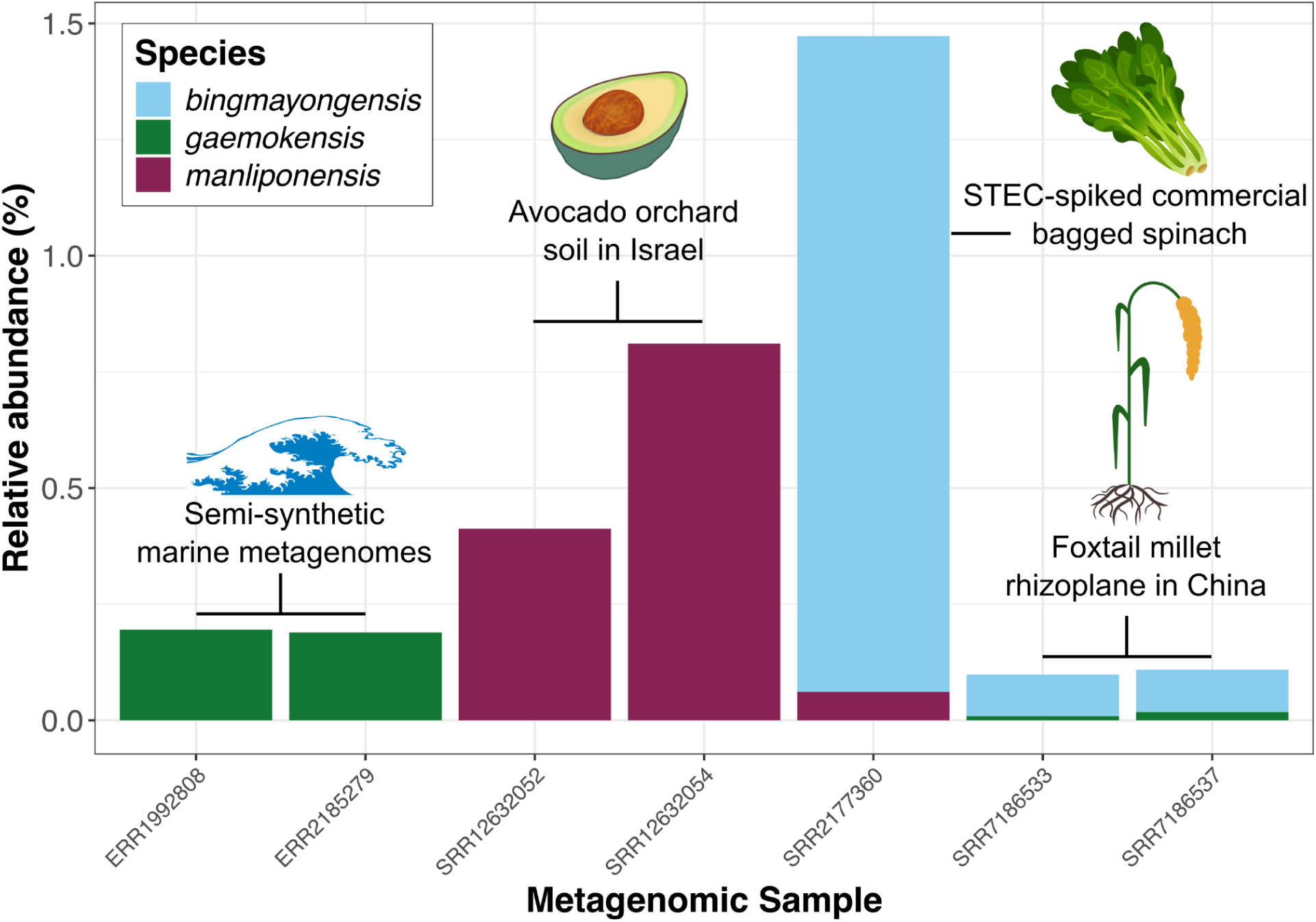
Rare *B. cereus* group species can be detected in metagenomes harboring species-specific markers. Relative abundance (percent, %; Y-axis) of rare *B. cereus* group species in public metagenomes harboring markers of rare *B. cereus* group species (i.e., using Pebblescout, a rare *B. cereus* group species *panC* allele was detected in the metagenomic sample with ≥80% coverage and/or ≥15% of all species-specific genes were detected in the sample with ≥80% coverage; X-axis).

In total, our MAG construction approach resulted in four high-quality *B. manliponensis* MAGs (CheckM contamination ≤5% and CheckM completeness ≥90%), all from one soil sample (NCBI SRA Run accession SRR12632054; **Supplementary Table S17**). Because both soil samples were biological replicates from the same orchard/location, collected on the same date [118], we selected one high-quality MAG to represent the *B. manliponensis* strain from this site (i.e., assembled with MEGAHIT and binned with SemiBin2; **Supplementary Table S17**). This high-quality MAG, which we refer to hereafter as the “Israeli avocado orchard” *B. manliponensis* strain, was used in subsequent steps.

Relative abundances were obtained using the mOTUs profiler. For readability, relative abundances of other species have been omitted (to view relative abundances of all species, see **Supplementary Figure S2**).

### South Korean *B. manliponensis* strains show a high degree of genomic diversity

We compared the Israeli avocado orchard *B. manliponensis* strain to all publicly available *B. manliponensis* genomes/MAGs (*n* = 3 genomes representing 2 unique strains): (i) JCM 15802 and CIP111019T, the original and re-sequenced genomes of the *B. manliponensis* type strain, which was isolated in 2011 from oil-contaminated foreshore tidal flat sediment of the Yellow Sea, South Korea (NCBI Assembly accessions GCA_000712595.1 and GCA_965139495.1, respectively; https://catalogue-crbip.pasteur.fr/fiche_catalogue.xhtml?crbip=CIP%20111019, accessed 12 July 2026) [21]; and (ii) jeju3-1, isolated in 2024 from coastal soil in Jeju, South Korea (NCBI Assembly accession GCA_041225435.1) [119].

The Israeli avocado orchard *B. manliponensis* strain was most closely related to the *B. manliponensis* type strain, sharing 99.60 ANI (via skani; **Figure 4A**) and differing by 8,057 SNPs (via Snippy; **Figure 4B**). Interestingly, South Korean strain jeju3-1 shared a relatively low degree of genomic similarity with both the South Korean *B. manliponensis* type strain and Israeli avocado orchard strain (95.05-95.39 ANI), despite its geographic origin, which was much closer to that of the type strain (**Figure 4AB**). Similar results were observed for accessory genomic content, as the South Korean *B. manliponensis* type strain was more similar to the Israeli avocado orchard strain than it was to the South Korean jeju3-1 strain (using Panaroo gene presence/absence and a Jaccard distance metric; **Figure 4C**, **Supplementary Figure S3**).

**Figure 4.**
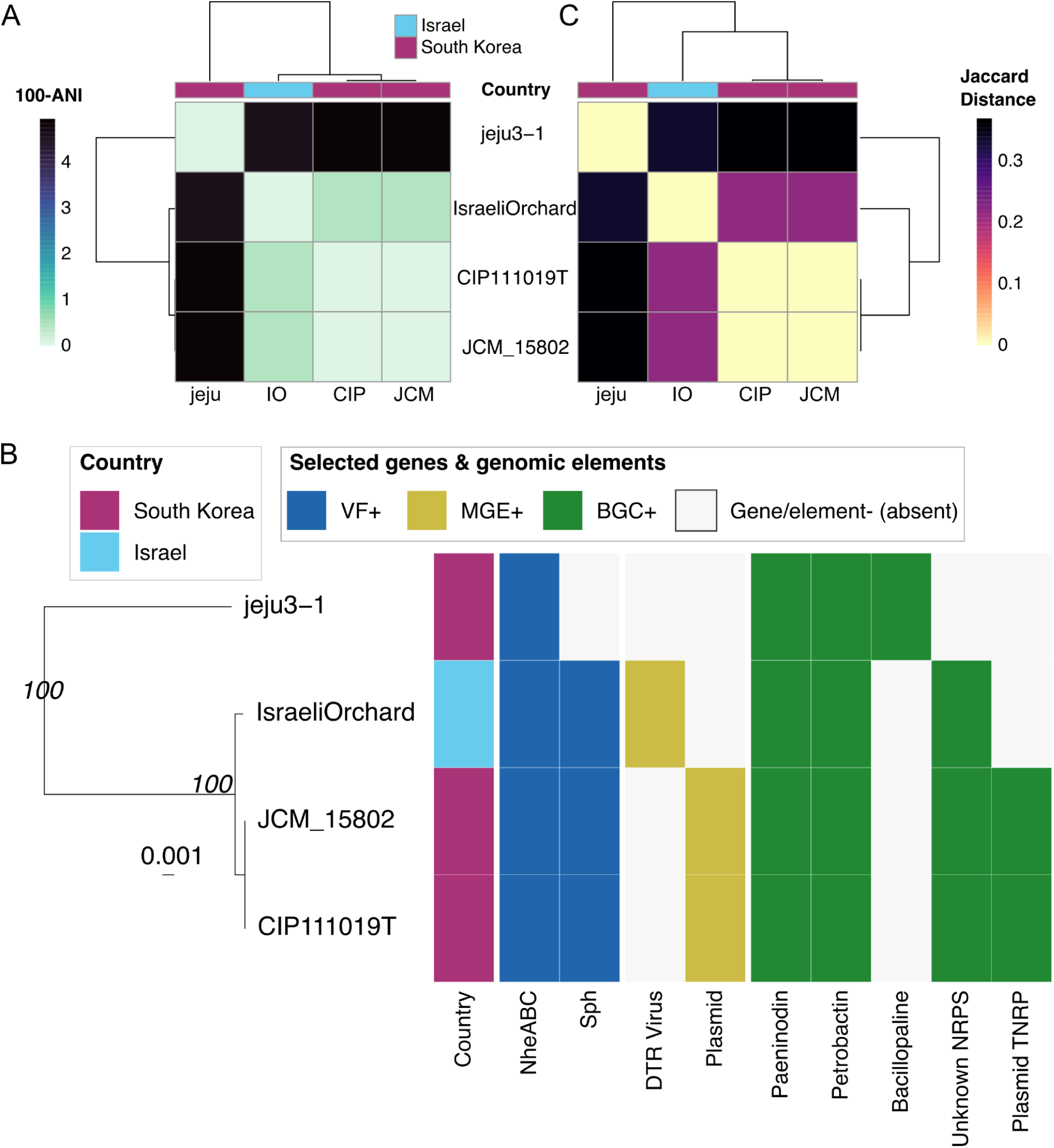
*B. manliponensis* from avocado orchard soil in Israel most closely resembles strain JCM 15802 from foreshore tidal flat sediment of the Yellow Sea. **(A)** Heatmap of pairwise average nucleotide identity (ANI) dissimilarity values (100-ANI), calculated between all *B. manliponensis* genomes/metagenome-assembled genomes (MAGs; *n* = 4) using skani. Dendrograms were produced via average linkage hierarchical clustering using ANI dissimilarities (100-ANI). **(B)** Maximum-likelihood phylogeny of all *B. manliponensis* genomes/MAGs. The heatmap to the right of the phylogeny shows (from left to right): (i) the country of origin for each genome/MAG (“Country”); (ii) presence and absence of selected virulence factors (“VF+”; detected using BTyper3’s implementation of tblastn); (iii) presence and absence of selected mobile genetic elements (“MGE+”; detected using geNomad); (iv) presence and absence of select biosynthetic gene clusters (“BGC+”; detected using antiSMASH and/or GECCO). The phylogeny was constructed using 684 core genes detected in all *B. manliponensis* genomes plus an outgroup (the *B. pseudomycoides* type strain). Branch support values (italicized) denote ultrafast bootstrap percentages (obtained using IQ-TREE). The phylogeny is rooted using the outgroup (omitted for readability), with branch lengths reported in substitutions per site. Metadata included in the heatmap was selected for readability (see **Supplementary Figures S3-S5** to view all data/datasets). Nhe, non-hemolytic toxin; Sph, sphingomyelinase C; DTR, direct terminal repeats; NRPS, nonribosomal peptide synthetase; TNRP, thioamide-nonribosomal peptide synthetase. An interactive version of this phylogeny can be viewed using BTracker (https://nextstrain.org/community/vigzy77/BTracker/Bacillus-cereus-group/BTyper3-sp-m <u>anliponensis(Core-genome-ML)</u>). **(C)** Heatmap of accessory genome dissimilarity for all *B. manliponensis* genomes/MAGs (*n* = 4). Heatmap shading shows Jaccard distances calculated between genomes/MAGs based on Panaroo gene presence/absene patterns. Dendrograms were produced via average linkage hierarchical clustering using Jaccard distances. For **(A)** and **(C),** the color strips above each heatmap show the country of origin for each genome/MAG.

Overall, our results indicate that (i) highly diverse *B. manliponensis* strains are circulating in South Korea; (ii) *B. manliponensis* can be detected in agricultural environments, in proximity to produce with the potential for human consumption; and (iii) *B. manliponensis* is not confined to a single geographic region, as the Israeli avocado orchard MAG constructed here represents the first genome of a *B. manliponensis* strain observed outside of South Korea.

### Homologs of virulence factor sphingomyelinase C are variably present within *B. manliponensis*

Using BTyper3’s implementation of translated nucleotide BLAST (tblastn) [107], no anthrax toxin genes (*cya, lef, pagA*) or cereulide synthetase (emetic toxin) core biosynthetic enzymes (*cesAB*) could be detected in any *B. manliponensis* genomes/MAGs (**Supplementary Tables S30-S33**), nor were they detected in any other MAGs from the *B. manliponensis*-harboring Israeli avocado orchard samples (i.e., SRR12632052 and SRR12632054, BTyper3 default parameters; **Supplementary Tables S39-S40**). However, all four *B. manliponensis* genomes/MAGs harbored homologs of non-hemolytic toxin (Nhe)-encoding *nheABC*, one of the major *B. cereus* group toxins involved in foodborne diarrheal toxicoinfection (**Figure 4B**, **Supplementary Tables S30-S33**) [1]. Interestingly, 3 of 4 (meta)genomes (i.e., all but jeju3-1) possessed a homolog of sphingomyelinase C-encoding *sph*, a *B. cereus* group virulence factor that has been shown to complement Nhe-induced cytotoxicity (**Figure 4B, Supplementary Tables S30-S33**) [120].

Altogether, *B. manliponensis* strains differ in virulence gene composition and thus may differ in cytotoxicity/toxin production, as is the case for other *B. cereus* group species [39]. However, future assessments of cytotoxicity and toxin production *in vitro* and *in vivo* will be needed for confirmation.

### Diverse viral sequences are detected in *B. manliponensis*, but only the type strain is predicted to harbor a plasmid

With regards to mobile genetic elements (MGEs), *B. manliponensis* strains each possessed their own unique viral sequence profile (**Figure 4B**, **Supplementary Figure S4**, **Supplementary Table S52**). Within the Israeli avocado orchard MAG specifically, a unique viral sequence with a direct terminal repeat (DTR) topology was detected (geNomad virus score = 0.9978; **Figure 4B**, **Supplementary Table S49**). When compared to all 356,776 prokaryotic prophage sequences within Prophage-DB [111], this DTR viral sequence belonged to a novel gene cluster family (i.e., it did not cluster with any Prophage-DB prophage sequences, nor any other predicted *B. manliponensis* viral sequences, based on protein content; **Figure 4B**, **Supplementary Table S52**). BLASTP (protein BLAST) of all viral proteins against NCBI’s nr (non-redundant protein sequence) database confirmed that nearly all individual proteins within this DTR viral sequence most closely resembled proteins from *B. cereus* group members (including other *B. manliponensis* strains; **Supplementary Table S56**), indicating that this unique viral sequence was likely not the result of an assembly/binning error.

While diverse viral sequences were detected in all *B. manliponensis* genomes, very few *B. manliponensis* contigs were predicted to be of plasmid origin. Specifically, the two *B. manliponensis* type strain genomes were each predicted to possess a single contig of plasmid origin (geNomad plasmid score >0.99 for both genomes), neither of which possessed conjugation genes (per geNomad; **Supplementary Table S48**). While the lack of plasmid-associated sequences in the Israeli avocado orchard *B. manliponensis* MAG could be a binning artifact, the jeju3-1 isolate genome (a complete genome, sequenced using long reads) also did not harbor plasmid-associated sequences (**Supplementary Table S48**). While these results indicate that *B. manliponensis* may harbor diverse viral sequences and relatively few plasmids, future *B. manliponensis* isolation and long-read WGS efforts are needed to accurately assess MGEs among *B. manliponensis*.

### The Israeli avocado orchard and *B. manliponensis* type strains possess a putative novel nonribosomal peptide synthetase

A total of 19 unique families of biosynthetic gene clusters (BGCs) were detected in *B. manliponensis* (**Supplementary Figure S5**, **Supplementary Table S54**), 3 of which contained previously characterized BGCs responsible for the production of known secondary metabolites (**Figure 4B**).

Specifically, all strains possessed BGCs that resembled BGCs responsible for the production of (i) paeninodin, a member of the lasso peptide class of ribosomally synthesized and post-translationally modified peptides (RiPPs) produced by *Paenibacillus dendritiformis* (MIBiG accession BGC0001356; **Supplementary Figure S6**) [121]; and (ii) petrobactin, a siderophore produced by *B. anthracis*, which is required for growth in macrophages and virulence in a mouse model (MIBiG accession BGC0000942; **Figure 4B, Supplementary Figure S7**) [122, 123]. The relatively distant jeju3-1 strain additionally harbored a BGC that closely resembled (iii) bacillopaline, a zincophore produced by *Paenibacillus mucilaginosus* (MIBiG accession BGC0002488; **Figure 4B, Supplementary Figure S8**) [124].

Notably, in the *B. manliponensis* type strain genomes and Israeli avocado orchard MAG, both antiSMASH and GECCO detected a putative novel nonribosomal peptide synthetase (NRP), which did not cluster with any MIBiG BGCs based on protein content (**Figure 4B, Supplementary Figure S9, Supplementary Table S54**). According to antiSMASH, this putative novel NRP most closely resembled NRPs xenotetrapeptide (MIBiG accession BGC0001132), gamexpeptide C (BGC0001128), pyreudione A-E (BGC0002075), and xenematide (BGC0001825), although it was relatively distant from all MIBiG BGCs (antiSMASH similarity score <0.75).

The lone contig in the *B. manliponensis* type strain genome (strain JCM 15802) predicted to be of plasmid origin (geNomad plasmid score = 0.9987) additionally harbored a BGC, which was detected by both antiSMASH and GECCO. Detected in both type strain genomes, this BGC did not cluster with any MIBiG BGCs based on protein content (**Figure 4B, Supplementary Table S54**). According to antiSMASH, this putative novel BGC was assigned to two classes (“thioamide-NRP” and “terpene-precursor”) and most closely resembled three NRPs (bicornutin A1 from *Xenorhabdus budapestensis*, MIBiG accession BGC0001135; xenematide from *Xenorhabdus nematophila AN6/1*, BGC0001825; and ririwpeptide ABC from *Photorhabdus laumondii subsp. laumondii TTO1*, BGC0002286), although with very low antiSMASH similarity scores (0.24-0.26; **Supplementary Figure S10**).

Taken together, our results suggest that (i) strains within *B. manliponensis* possess BGCs that resemble those responsible for the production of known secondary metabolites (i.e., paeninodin, petrobactin, and bacillopaline); (ii) even with few genomes/MAGs available, *B. manliponensis* harbors putative novel BGCs (e.g., the NRP identified here), some of which (iii) are harbored on plasmids (e.g., the thioamide-NRP/terpene-precursor BGC identified here), despite *B. manliponensis* harboring few contigs of predicted plasmid origin.

### *B. bingmayongensis* is present in the rhizoplane of Chinese foxtail millet and an STEC-spiked commercial spinach sample

While *B. bingmayongensis*-specific genes were detected in 134 metagenomic samples, only two samples (NCBI SRA Run accessions SRR7186533 and SRR7186537), both collected September 2013 from the rhizoplane of *Setaria italica* (foxtail millet) in China (Shanxi, Yangling), possessed >15% of all *B. bingmayongensis*-specific genes (≥80% coverage per Pebblescout; **Figure 2B**, **Supplementary Table S3**) [125]. Only one additional sample (NCBI SRA Run accession SRR2177360), from commercial bagged spinach purchased at a grocery store and subsequently spiked with Shiga toxin-producing *Escherichia coli* (STEC), harbored the *B. bingmayongensis panC* allele (85.71% coverage); this sample additionally harbored 18/195 (9.23%) *B. bingmayongensis*-specific genes (≥80% coverage per Pebblescout; **Figure 2B**, **Supplementary Table S7**) [126].

Using the mOTUs profiler, *B. bingmayongensis* was present in the two Chinese foxtail millet rhizoplane samples with low relative abundance (<0.10%; **Figure 3**, **Supplementary Tables S12-S13**). However, in the STEC-spiked commercial spinach, *B. bingmayongensis* had the highest relative abundance of any rare *B. cereus* group species queried in this study (**Figure 3, Supplementary Table S11**). Further, additional rare *B. cereus* group species were detected in all three *B. bingmayongensis*-harboring samples (i.e., *B. gaemokensis* in both Chinese foxtail millet rhizoplane samples, and *B. manliponensis* in the STEC-spiked commercial spinach sample), albeit with very low relative abundance (**Figure 3**, **Supplementary Tables S11-S13**). In line with this, no *B. gaemokensis* or *B. manliponensis* MAGs were recovered from these samples.

MAG construction efforts for the two Chinese foxtail millet rhizoplane samples (SRR7186533 and SRR7186537) resulted in a collection of *B. cereus* group MAGs with low completeness and high contamination (per CheckM), none of which could reliably be assigned to *B. bingmayongensis*. The quality of *B. cereus* group MAGs from the STEC-spiked commercial bagged spinach sample (SRR2177360) was higher, and 11 of the resulting MAGs shared >97 ANI with the *B. bingmayongensis* type strain genome (**Supplementary Table S18**). Of these, 10 MAGs either (i) harbored the *B. bingmayongensis panC* allele and/or (ii) were assigned to *B. bingmayongensis* using GTDB-Tk, and were thus considered to be *B. bingmayongensis* MAGs (**Supplementary Table S18**). While all *B. bingmayongensis* MAGs were incomplete (CheckM Completeness <70%), we obtained a 1.7 Mb MAG with low contamination (<5% per CheckM), which was classified as *B. bingmayongensis* using both skani and GTDB-Tk, and possessed a full-length *B. bingmayongensis panC* allele (100% coverage and 99.4% nucleotide identity using BTyper3; **Supplementary Table S18**). This MAG, which was assembled with MEGAHIT, binned with SemiBin2, and is referred to hereafter as the “commercial spinach” *B. bingmayongensis* strain, was selected to represent *B. bingmayongensis* from STEC-spiked commercial bagged spinach sample SRR2177360 (**Supplementary Table S18**).

### Multiple *B. bingmayongensis* lineages are circulating in China

We compared the commercial spinach *B. bingmayongensis* strain to all publicly available *B. bingmayongensis* genomes/MAGs (*n* = 4 genomes representing 4 unique strains): (i) FJAT-13831, the *B. bingmayongensis* type strain, which was isolated in 2010 from a soil sample collected from the pit of Emperor Qin’s Terra-cotta Warriors in Shaanxi, China (NCBI Assembly accession GCA_000299035.1) [10]; (ii) P86, isolated in 2016 from soil in Zhengzhou, China (NCBI Assembly accession GCA_019753385.1) [127]; (iii) BUPM19, isolated in 1994 from soil in Tunisia (NCBI Assembly accession GCA_034427855.1) [128]; (iv) XF8, isolated in 2020 from peanut rhizosphere soil in Henan, China (NCBI Assembly accession GCA_017498755.1). Using ANI and phylogenetic methods, the commercial spinach *B. bingmayongensis* strain was most closely related to Chinese peanut rhizosphere soil strain XF8 (98.93 ANI via skani; **Figure 5**). The remaining three *B. bingmayongensis* genomes (two from China and one from Tunisia) formed a separate clade within the *B. bingmayongensis* phylogeny (ultrafast bootstrap support = 100%; **Figure 5**).

**Figure 5.**
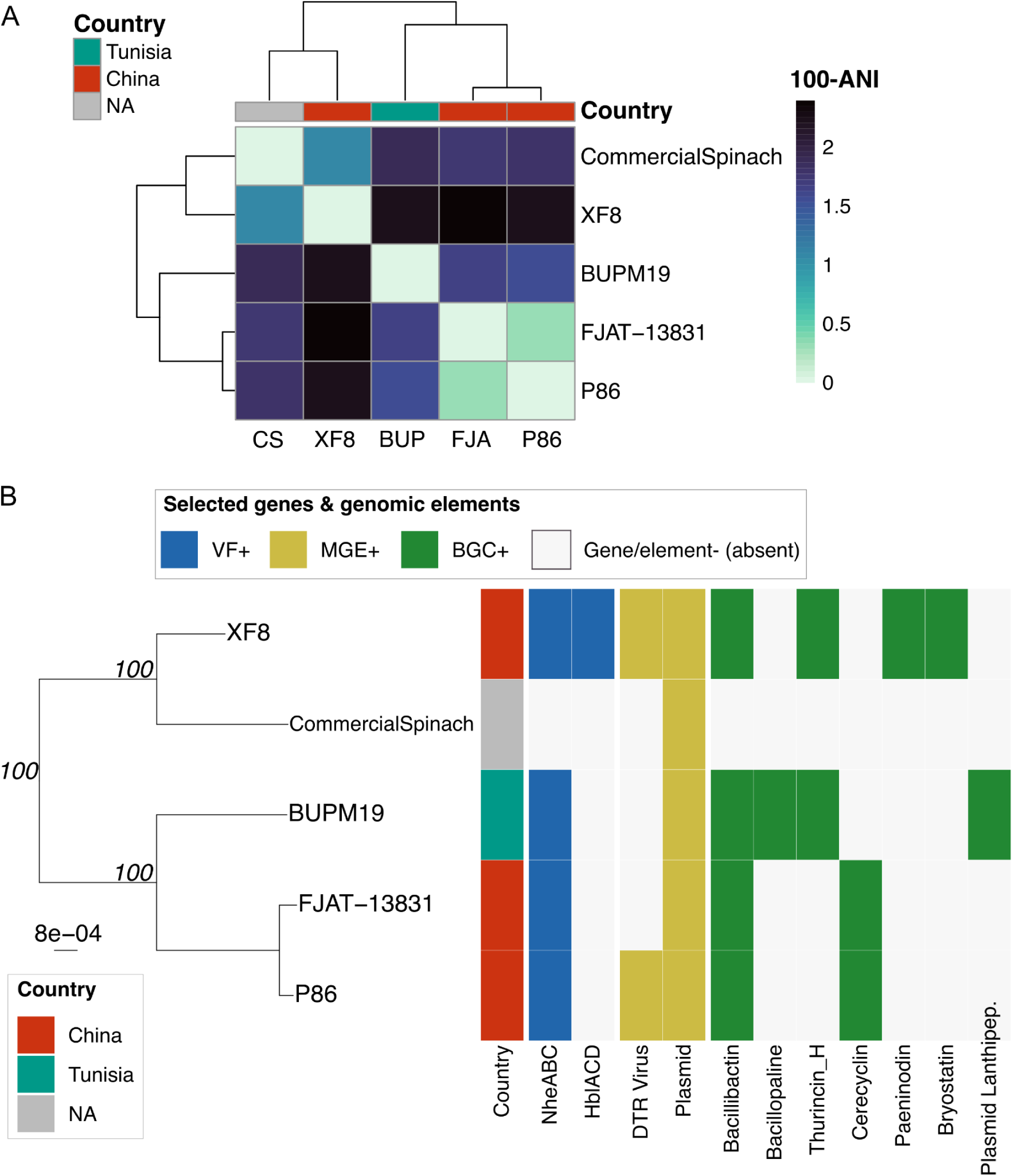
*B. bingmayongensis* from STEC-spiked commercial spinach most closely resembles strain XF8 from peanut rhizosphere soil in China. (A) Heatmap of core genome dissimilarity for all *B. bingmayongensis* genomes/metagenome-assembled genomes (MAGs; *n* = 5). Heatmap shading shows pairwise average nucleotide identity (ANI) dissimilarity values (100-ANI; calculated via skani). The color strip above the heatmap shows the country of origin for each genome/MAG. Dendrograms were produced via average linkage hierarchical clustering using ANI dissimilarities (100-ANI). **(B)** Maximum-likelihood phylogeny of all *B. bingmayongensis* genomes/MAGs. The heatmap to the right of the phylogeny shows (from left to right): (i) the country of origin for each genome/MAG (“Country”); (ii) presence and absence of selected virulence factors (“VF+”; detected using BTyper3’s implementation of tblastn); (iii) presence and absence of selected mobile genetic elements (“MGE+”; detected using geNomad); (iv) presence and absence of select biosynthetic gene clusters (“BGC+”; detected using antiSMASH and/or GECCO). The phylogeny was constructed using 882 core genes detected in all *B. bingmayongensis* genomes plus an outgroup (the *B. pseudomycoides* type strain). Branch support values (italicized) denote ultrafast bootstrap percentages (obtained using IQ-TREE). The phylogeny is rooted using the outgroup (omitted for readability), with branch lengths reported in substitutions per site. Metadata included in the heatmap was selected for readability (see **Supplementary Figures S11-S14** to view all data/datasets). NA, not available; Nhe, non-hemolytic toxin; Hbl, hemolysin BL; DTR, direct terminal repeats; Lanthipep., lanthipeptide. An interactive version of this phylogeny can be viewed using BTracker (https://nextstrain.org/community/vigzy77/BTracker/Bacillus-cereus-group/BTyper3-sp-bingmayongensis(Core-genome-ML)).

Overall, our results show that (i) multiple *B. bingmayongensis* lineages are circulating in China, as some Chinese strains are more closely related to Tunisian soil strain BUPM19 than to other Chinese strains; and (ii) as observed for *B. manliponensis*, *B. bingmayongensis* can be detected in agricultural environments, in proximity to foods with the potential for human consumption.

### Hemolysin BL homologs are variably present within *B. bingmayongensis*

Anthrax toxin genes (*cya*, *lef*, *pagA*) and cereulide (emetic toxin) synthetase core biosynthetic enzymes (*cesAB*) were not detected in any *B. bingmayongensis* genomes/MAGs (**Supplementary Tables S34-S38**), nor were they detected in any other MAGs from *B. bingmayongensis*-harboring samples (i.e., Chinese foxtail millet rhizoplane samples SRR7186533 and SRR7186537, and STEC-spiked commercial bagged spinach sample SRR2177360; BTyper3 default parameters, **Supplementary Tables S41-S43**). As observed for *B. manliponensis*, genes encoding non-hemolytic toxin (*nheABC*) could be detected in all *B. bingmayongensis* genomes except the commercial spinach MAG, likely due to the MAG being incomplete (CheckM Completeness = 33.24%; **Figure 5, Supplementary Table S20**). Interestingly, homologs of hemolysin BL (*hblACD*), a *B. cereus* group toxin involved in foodborne diarrheal toxicoinfection [1], were detected in strain XF8 from peanut rhizosphere soil in China, but were absent from all other *B. bingmayongensis* genomes/MAGs (**Figure 5, Supplementary Tables S20, S34-S38**). Thus, like *B. manliponensis*, virulence factor heterogeneity can be observed in *B. bingmayongensis*, and our results highlight the need for future strain-specific assessments of *B. bingmayongensis* cytotoxicity/toxin production.

### All *B. bingmayongensis* genomes and MAGs are predicted to harbor plasmids

With regards to MGEs, all *B. bingmayongensis* genomes possessed at least one contig predicted to be of viral origin (per geNomad), with three genomes (strains XF8, BUPM19, and FJAT-13831) harboring predicted prophage (**Supplementary Figure S13, Supplementary Table S51**). In two genomes (strains XF8 and P86), two viral contigs were assigned to the same GCF (gene cluster family) based on their protein composition (**Figure 5B, Supplementary Table S53**). No publicly available prophage sequences (from Prophage-DB) were detected within this family, highlighting its novelty, and in strain P86, the viral sequence possessed direct terminal repeats (DTR; **Figure 5B**, **Supplementary Figure S15, Supplementary Table S53**).

Unlike *B. manliponensis*, where a single unique strain (the type strain) was predicted to harbor a plasmid, all *B. bingmayongensis* genomes/MAGs possessed at least one contig predicted to be of plasmid origin (**Figure 5B, Supplementary Table S50**). Further, in all but 1 genome (BUPM19, the Tunisian strain), at least one plasmid-associated contig harbored a MOB conjugation element, indicating potential for plasmid mobility (per geNomad; **Supplementary Table S50**). These results indicate that plasmids appear to be an important component of the *B. bingmayongensis* genome, although future isolation and long-read WGS efforts are needed to confirm this.

### *B. bingmayongensis* strain BUPM19 harbors a putative novel cyclic lactone autoinducer / class II lanthipeptide BGC on a plasmid

A total of 37 unique families of BGCs were detected in the 5 *B. bingmayongensis* genomes/MAGs (**Supplementary Figure S14**), including 6 that contained previously characterized BGCs responsible for the production of known secondary metabolites (**Figure 5B, Supplementary Table S55**). All genomes but the incomplete commercial spinach MAG possessed a BGC, which was similar to the BGC responsible for the production of siderophore bacillibactin (MIBiG accession BGC0002695; **Figure 5B**, **Supplementary Figure S16**) [129]. Other BGCs resembling those responsible for the production of other known secondary metabolites (e.g., zincophore bacillopaline, MIBiG BGC0002488; bacteriocin thurincin H, MIBiG BGC0000600; bacterocin cerecyclin, MIBiG BGC0002447; lasso peptide paeninodin, MIBiG BGC0001356; macrolide lactone/protein kinase C modulator bryostatin, MIBiG BGC0000205) were variably present among strains (**Figure 5B, Supplementary Figures S17-S21**) [130–132].

Notably, both antiSMASH and GECCO detected a BGC exclusive to strain BUPM19, which did not cluster with any MIBiG BGCs and was located on a predicted plasmid (geNomad plasmid score = 0.9854; **Figure 5B, Supplementary Table S50**). Classified by antiSMASH as both a “cyclic-lactone-autoinducer” and “lanthipeptide-class-ii” BGC, this BGC most closely resembled cytolysin ClyLl, a RiPP from plasmid pAD1 (MIBiG BGC0000504, antiSMASH similarity score 0.81; **Figure 5B, Supplementary Figure S22**). Altogether, these findings suggest that *B. bingmayongensis* is capable of producing a diverse array of known and novel secondary metabolites, some of which are predicted to be harbored on plasmids.

### *B. gaemokensis* can be detected in semi-synthetic marine metagenomes with low relative abundance

*B. gaemokensis*-specific genes were extremely rare in SRA, with only 6 samples harboring one or more gene(s) at ≥80% coverage (**Figure 2B, Supplementary Table S4**). However, the *B. gaemokensis panC* allele was additionally detected in two of these samples (92.1% coverage per Pebblescout), both submitted to NCBI as “semi-synthetic marine metagenomes”, which had been created for benchmarking metagenomic tools/pipelines (NCBI SRA Run accessions ERR1992808 and ERR2185279; **Supplementary Table S8**). Using the mOTUs profiler, *B. gaemokensis* was detected in both samples, but with low relative abundance (<0.20%; **Figure 3**, **Supplementary Tables S14-S15**). While MAG construction resulted in a total of 6 MAGs sharing >99.0 ANI with the *B. gaemokensis* type strain genome (3 from ERR1992808 and 3 from ERR2185279), all were of low quality (CheckM Completeness <30% or Contamination >100%) and could not be reliably assigned to *B. gaemokensis* using either GTDB-Tk or BTyper3. Anthrax toxin genes (*cya*, *lef*, *pagA*) and cereulide (emetic toxin) synthetase core biosynthetic enzymes (*cesAB*) were not detected in any MAGs constructed from the *B. gaemokensis*-harboring samples (**Supplementary Tables S44-S45**).

## DISCUSSION

### Rare bacterial species: a sampling problem, or a biological/ecological reality?

Members of the *B. cereus* group are considered to be virtually ubiquitous in nature [1], and as of 2026, nearly 15,000 genomes derived from *B. cereus* group isolates are available in public databases [59]. However, not all species are equally represented, as some species (e.g., *B. anthracis*, *B. cereus s.s.*) are represented by thousands of genomes, while others are represented by fewer than five [59]. Further, while the *B. cereus* group can be partitioned into 18 genomospecies (delineated using a 92.5 ANI threshold as described previously) [37], eight of these genomospecies (44.4%) account for >98% of all publicly available *B. cereus* group genomes (per BTyperDB, accessed September 2025) [59]. Here, we searched all of NCBI SRA for metagenomic samples harboring the rarest of the *B. cereus* group species–some represented by just a single genome, despite having been sequenced/discovered over a decade ago–and we identify multiple metagenomic samples harboring three longstanding rare species (i.e., *B. manliponensis*, *B. bingmayongensis*, and *B. gaemokensis*).

There are multiple possible explanations as to why some *B. cereus* group species are represented by very few genomes in public databases, the first being that these species are, indeed, truly “rare” in nature (i.e., present with low abundance, possibly confined to a specific geographic location/environment, due to e.g., lower competitive ability and/or a narrow ecological niche) [133, 134]. It is possible that all (or a subset of) the four rare *B. cereus* group species queried here are restricted to a narrow range of environments/conditions for survival and growth (e.g., a narrow temperature range, highly specific nutrient requirements) [134]. Alternatively/additionally, these rare species may struggle to out-compete other species, including other members of the *B. cereus* group [134]. Interestingly, we observed that 5 of the 7 rare-species harboring metagenomic samples queried here (i.e., all but the semi-synthetic marine metagenomes) also harbored other *B. cereus* group species (**Supplementary Figure S2, Supplementary Tables S9-S15**). Thus, future studies assessing growth and cooperation/competition among diverse *B. cereus* group strains (including rare species) under different environmental conditions are needed, as our results indicate that multiple *B. cereus* group species may frequently occupy the same ecological niche.

While rarity could indeed be a reality, sampling/sequencing biases could also explain why rare *B. cereus* group species are under-represented in public databases. Some bacterial species are endemic to particular geographical regions (e.g., *B. anthracis*) [135], and it is possible that current sampling, isolation, and/or sequencing efforts are not concentrated in world regions where rare *B. cereus* group species are most commonly found [136, 137]. Alternatively/additionally, commonly used isolation protocols for *B. cereus* group species may unintentionally select against rare *B. cereus* group species. For example, a recent study showed that incubation at 50°C (i.e., a temperature 15-20°C higher than that recommended by the U.S. Food and Drug Administration Bacteriological Analytical Manual [FDA BAM] for enumeration and confirmation of *B. cereus* group strains in food) [138] greatly increased the number of food samples testing positive for *B. cytotoxicus* [139]. Similarly, choice of agar can select for/against different *Bacillus* strains [140]. While the metagenomic approach used here bypasses isolation biases, it is further possible that metagenomic sample processing and/or DNA extraction protocols could be biased against rare *B. cereus* group species, as some protocols may not effectively capture Gram-positive sporeformers [141, 142]. Thus, continued expansion of metagenomic sequencing efforts worldwide, using protocols capable of capturing Gram-positive sporeformers, will likely continue to uncover novel genomes of rare *B. cereus* group species.

### Rare *B. cereus* group species are present in food and agricultural environments

Despite having been published and/or first sequenced over a decade ago, the four rare *B. cereus* group species are absent from many clinically and industrially important microbiological resources (e.g., the FDA BAM) [138], and their growth (or lack thereof) has not been evaluated using many popular isolation protocols (e.g., standard temperatures and media used for *B. cereus* group isolation) [138, 143]. This is likely largely due to: (i) their rarity, as prior to 2021, all rare species were represented by a single strain; and (ii) the fact that, until recently, all rare species strains had been isolated from environments with limited potential for human consumption/contact (e.g., prior to 2021, tidal flat sediment from the Yellow Sea, pit soil from Emperor Qin’s Terra-cotta warriors in China, and soil in Papua New Guinea) [10, 14, 18, 21].

Here, we show that rare *B. cereus* group species can be detected in agricultural environments and foods with the potential for human consumption, including raw spinach being sold commercially (*B. bingmayongensis*), the rhizoplane/rhizosphere of foxtail millet and peanut plants in China (*B. bingmayongensis*), and soil from a freshwater-irrigated avocado orchard in Israel (*B. manliponensis*). Further, we show that *B. maniponensis* and *B. bingmayongensis* possess homologs of known virulence factors, including non-hemolytic toxin (Nhe; all complete genomes), hemolysin BL (Hbl; 1 *B. bingmayongensis* strain), and sphingomyelinase C (Sph; 3 *B. manliponensis* genomes; **Figures 4-5**). It is important to note that merely detecting a virulence factor-harboring *B. cereus* group strain in a food sample does not mean that particular strain represents a food safety or public health threat. Foodborne illnesses caused by *B. cereus* group members are complex and governed by a range of interplaying factors (e.g., strain, food matrix, temperature, pH, oxygen, infectious dose) [1, 39, 144], and virtually all *B. cereus* group genomes possess virulence factor homologs (e.g., Nhe) [38]. However, our results highlight the need for studies assessing food safety and public health risks associated with rare *B. cereus* group species (via e.g., studies assessing toxin production, cytotoxicity, growth, and survival in diverse matrices, under clinically and industrially relevant conditions).

### Unassembled metagenomes are a potential treasure trove of rare species

In the analysis of shotgun metagenomic data, taxonomic profiling, *de novo* assembly, and binning are computationally intensive and time-consuming processes. While massive ongoing MAG generation efforts (e.g., via mOTUs, SPIRE) [88, 89] are capturing more microbial diversity than ever before, all have thus far failed to generate any MAGs from rare *B. cereus* group species. Here, we developed a targeted approach for rapidly identifying putative rare species-harboring metagenomes in NCBI’s SRA database in seconds; prioritized samples can then undergo taxonomic profiling, and if desired, *de novo* assembly and binning.

While our method rapidly identified rare species in unassembled metagenomes, it is important to point out its limitations. First, while Pebblescout searches themselves take only seconds (or less), our method requires the identification of rare species markers (i.e., species-specific genes or alleles) as a prerequisite–a step that, for some taxa/datasets, can be time-consuming and computationally expensive (due to the computational demands of e.g., ortholog identification, multiple sequence alignment, phylogeny construction). Second, as a proof-of-concept, we focused our efforts solely on metagenomic samples with a high likelihood of harboring rare *B. cereus* group species. However, hundreds of samples possessed a smaller selection of species-specific markers (**Figure 2B**). While many of these samples are likely false positives (e.g., they possess a single species-specific gene, which was acquired/disseminated via horizontal gene transfer; a species-specific gene was a contaminant within the original rare *B. cereus* group species genomes used for marker selection), it is likely that our conservative selection criteria missed rare species-harboring samples and was biased towards strains more closely resembling rare species type strains. Third, the *de novo* assembly and binning process can introduce contamination into MAGs, which can affect our downstream pan-genomic analyses [145]. While we attempted to combat this by testing multiple assembly/binning method combinations and employing strict quality control thresholds, we encourage readers to be aware of the limitations of metagenomics. Finally, linking extrachromosomal elements (e.g., plasmids) to individual MAGs is challenging, and sometimes impossible, for current binning methods [146]. This limitation is particularly problematic for assessing *B. cereus* group virulence potential, as some of the most critically important virulence factors from a public health perspective (e.g., anthrax toxin– and cereulide synthetase-encoding genes) are harbored on plasmids [1]. To combat this, we evaluated anthrax and cereulide synthetase gene presence/absence at the whole-metagenome level, in addition to the individual MAG level, and we encourage future users of our method to do the same.

Overall, our efforts posit unassembled metagenomes in public databases as a potential treasure trove of rare species genomes, which can be used to augment isolate genome collections. We anticipate that the targeted approach developed here can be used for the identification of other rare species, as well as novel species discovery.

## Acknowledgments

Computation was enabled by resources provided by (i) High Performance Computing Center North (HPC2N; Umeå University, Umeå, Sweden), and (ii) the National Academic Infrastructure for Supercomputing in Sweden (NAISS), partially funded by the Swedish Research Council through grant agreement no. 2022-06725. Images used within Figure 3 are within the public domain and are available via https://publicdomainvectors.org/en/free-clipart/Wavy-vector-graphics/1677.html, https://publicdomainvectors.org/en/free-clipart/Avocado-cut-in-half-vector-clip-art/18093.html, and https://publicdomainvectors.org/en/free-clipart/Spinach/44545.html.

## Author contributions

Computational analyses were performed by all co-authors. V Raghuram conceptualized the study, which was funded by LMC. LMC wrote the manuscript with input from all co-authors.

