## Supplementary Text for "Hiding in plain sight: rare *Bacillus cereus* group species are present among unassembled metagenomic samples in public databases"

**Identification of validly published and effective *B. cereus* group species.** A comprehensive list of validly published and effective *B. cereus* group species (as of 28 June 2026) was obtained in four major steps (Steps 1-4). First, in Step 1, we obtained a list of all validly published and effective *B. cereus* group species published through 2020 (i.e., Figure 1 of our 2022 *B. cereus* group taxonomy review) [1]; we then cross-referenced this initial list of *B. cereus* group species with all species assigned to the genus "*Bacillus*" in the List of Prokaryotic names with Standing in Nomenclature (LPSN; https://lpsn.dsmz.de/genus/bacillus, accessed 28 June 2026) [2, 3]. Species listed as "validly published under the ICNP" (International Code of Nomenclature of Prokaryotes) [4] in the "Nomenclatural status" column were considered to be "validly published" *B. cereus* group species, with the following two exceptions/clarifications: (i) one species (*B. weihenstephanensis*) was classified as "validly published under the ICNP", but was listed as a "synonym" in the "Taxonomic status" column [5]. Due to the continued importance of *B. weihenstephanensis* in the food industry [1], we classified *B. weihenstephanensis* as a validly published species, but noted that it is now considered a later heterotypic synonym of *B. mycoides* (per the ICNP/LPSN). (ii) After the publication of our *B. cereus* group taxonomy review [1], three of the four original effective *B. cereus* group species (i.e., "*B. bingmayongensis*", "*B. gaemokensis*", and "*B. manliponensis*") were reclassified as "validly published" species per the ICNP [6, 7]. Altogether, Step 1 resulted in an initial list of 22 validly published *B. cereus* group species (i.e., *B. albus*, *B. anthracis*, *B. bingmayongensis*, *B. cereus s.s.*, *B. cytotoxicus, B. fungorum, B. gaemokensis*, *B. luti, B. manliponensis*, *B. mobilis, B. mycoides, B. nitratireducens, B. pacificus, B. paramycoides, B. paranthracis, B. proteolyticus, B. pseudomycoides, B. thuringiensis, B. toyonensis*, *B. tropicus, B. weihenstephanensis, B. wiedmannii*) and 1 "effective" species (i.e., "*B. clarus*").

After constructing an initial list of validly published and effective *B. cereus* group species (i.e., those published through 2020), we added additional *B. cereus* group species published from 2021 onward by cross-referencing the LPSN list of *Bacillus* species (described above) with the National Center for Biotechnology Information (NCBI) Taxonomy database (Step 2) [8] and the Genome Taxonomy Database (GTDB; Step 3) [9]. For Step 2, all species listed as members of the "*Bacillus cereus* group" in NCBI's Taxonomy database (https://www.ncbi.nlm.nih.gov/Taxonomy/Browser/wwwtax.cgi?id=86661, accessed 28 June 2026) [8] were cross-referenced with the LPSN as described above. Species classified in the LPSN as "validly published under the ICNP" were considered to be validly published species, and species that were listed in the LPSN but were not classified as "validly published under the ICNP" were considered to be effective species, with the following clarifications/exceptions: (i) one species (*B. hominis*) was classified as "validly published under the ICNP", but was listed as a "synonym" in the "Taxonomic status" column [10]. Mirroring our approach for *B. weihenstephanensis* (described above), we classified *B. hominis* as a validly published species, but noted that it is now considered a later heterotypic synonym of *B. mycoides* (per the ICNP/LPSN). (ii) Three "species" names listed in NCBI Taxonomy were not present in the LPSN and were thus not included in our list of *B. cereus* group species, i.e., "*B. bombysepticus*", a species name used by GTDB, which is not commonly used by the *B. cereus* group community (supporting this: as of 28 June 2026, there are 45 NCBI BioSamples associated with NCBI's "*B. bombysepticus*" taxon, with the earliest two [SAMN02117259 and SAMN17068873, submitted in 2013 and 2020, respectively] having no linked sequencing data whatsoever, and the remainder submitted to NCBI from 2021 onward, after GTDB/GTDB-Tk were published in 2018/2020) [11–13]; "*B.* cf. *cereus* V4.BE.30", which was only represented in NCBI by a partial 16S rDNA gene (NCBI Nucleotide accession AJ244688.1) and had zero genomes available; and "*B. samanii*", also represented in NCBI by partial gene sequences, with zero genomes available (i.e., 13 NCBI Nucleotide accessions, 12 of which are partial 16S rDNA sequences; https://www.ncbi.nlm.nih.gov/nuccore/?term=txid411155[Organism:noexp], accessed 28 June 2026). Overall, this search of NCBI resulted in an additional five validly published *B. cereus* group species (i.e., *B. basilensis, B. dicomae, B. hominis, B. paramobilis, B. sanguinis*) and three effective *B. cereus* group species (i.e., "*B. altaicus*", "*B. shihchuchen*", "*B. silvicola*").

In Step 3, *B. cereus* group species from the latest version of GTDB (v232) [9] were cross-referenced against LPSN to identify additional validly published and effective *B. cereus* group species (i.e., those published after 2020, which were not considered to be part of NCBI's *B. cereus* group taxon). Briefly, all unique named species assigned to GTDB's "*Bacillus_A*" genus (i.e., a monophyletic clade within the GTDB phylogeny containing all known *B. cereus* group members) were cross-referenced against LPSN as described above. This search of GTDB resulted in five additional validly published *B. cereus* group species (i.e., *B. arachidis, B. bruguierae, B. multifaciens, B. rhizoplanae, B. yunxiaonensis*) and one effective *B. cereus* group species (i.e., "*B. pfraonensis*").

During our search, we noticed that NCBI's "*B. cereus* group" taxon was missing several validly published and effective species as child taxa (e.g., *B. bingmayongensis*, *B. fungorum*) [14, 15]. While these species were present in GTDB, other published *B. cereus* group species (e.g., *B. pacificus*, *B. weihenstephanensis*) were absent from GTDB (i.e., GTDB merges them with other *B. cereus* group species, as we have discussed previously) [1]. We thus searched for potential *B. cereus* group species in the LPSN, which met the following three conditions (Step 4): (i) the species was not listed as a child taxon of NCBI's *B. cereus* group taxon (per Step 2); (ii) the species was part of GTDB's *B. cereus* group (i.e., assigned to GTDB's "*Bacillus_A*" genus), but was not included as a GTDB species (due to e.g., GTDB's merging practices; per Step 3); (iii) the species was published after 2019 (as our previous review from 2022 included all validly published and effective species published through 2020, per Step 1) [1]. To accomplish this, the LPSN website (https://lpsn.dsmz.de/genus/bacillus, accessed 28

June 2026) was used to identify all publications describing novel/amended *Bacillus* species published from 2020 (the final year included in our *B. cereus* group taxonomic review) [1] to the present (i.e., 28 June 2026). For each novel/amended *Bacillus* species, the type strain genome's NCBI accession number (e.g., a NCBI Nucleotide accession number) was acquired from the corresponding publication and supplied to the NCBI website search function (https://www.ncbi.nlm.nih.gov/, accessed 28-29 June 2026), with the following exceptions: (i) the papers reportedly describing "*B. arbutinivorans*", "*B. cheonanensis*", and "*B. dabaoshanensis*" did not provide any NCBI accession numbers linked to whole genomes [16–18], and NCBI itself had no genomes linked to the taxa (only 16S sequences as of 29 July 2026); (ii) for "*B. oceanicus*" [19], no prokaryotic species with the name "*oceanicus*" were present in NCBI's Taxonomy database or GTDB (accessed 29 June 2026). The suffix of the NCBI GenBank Assembly accession linked to the type strain genome accession (i.e., the unique genome identifier, excluding the prefix "GCA_") was then searched in GTDB v232; if the type strain belonged to GTDB's "*Bacillus_A*" genus, the species was considered to be a member of the *B. cereus* group. If the type strain genome was not present in GTDB, the type strain (or species representative genome, if a type strain was not assigned in NCBI) was downloaded and supplied as input to GTDB-Tk v2.3.2 (i.e., the "classify_wf" workflow with GTDB database version r214 and default settings) [20]. This final effort yielded one additional effective *B. cereus* group species (i.e., "*B. pretiosus*").

**Calculation of average nucleotide identity values between species type strain genomes.** Average nucleotide identity (ANI) values were calculated between all species type strain genomes (*n* = 38; **Supplementary Table S1**) using FastANI v1.33 [21] and default settings (see section "Identification of validly published and effective *B. cereus* group species" above).

12. **Parks DH, Chuvochina M, Waite DW, Rinke C, Skarshewski A, *et al.*** A

standardized bacterial taxonomy based on genome phylogeny substantially revises the tree of life. *Nat Biotechnol* 2018;36:996–1004.
